# Blebs regulate phosphoinositides distribution and promote cell survival through the Septin-SH3KBP1-PI3K axis

**DOI:** 10.1101/2024.07.09.602823

**Authors:** Zifeng Zhen, Chunlei Zhang, Jiayi Li, Congying Wu

## Abstract

Cells rely on substrate adhesion to activate diverse signaling pathways essential for survival. In the absence of proper adhesion, cells undergo anoikis, a form of programmed cell death. Poorly attached cells often exhibit rounded morphologies and form small round protrusions called blebs. While the role of blebs in amoeboid migration is well-documented, recent studies have highlighted their function in anoikis resistance with the help of curvature-recognized proteins. However, little is known about whether the most abundant membrane components-phospholipids function in blebs-facilitated cell survival. Here, we found enriched PI(3,4,5)P3 and diluted PI(4,5)P2 at the bleb membrane, compared to non-bleb plasma membrane regions. Our experimental results showed that both phosphoinositides (PIs) have restricted diffusion pattern between the bleb and non-bleb membranes. Subsequently, we reveal SH3KBP1 to play a crucial role in recruiting PI3K and organizing the Septin-SH3KBP1-PI3K complex at the bleb necks. This process then contributes to differential PIs distribution and anoikis resistance. These novel insights into PIs dynamics elucidate their role in cell blebbing, cancer cell behavior and metastasis, presenting potential targets for new therapeutic interventions.

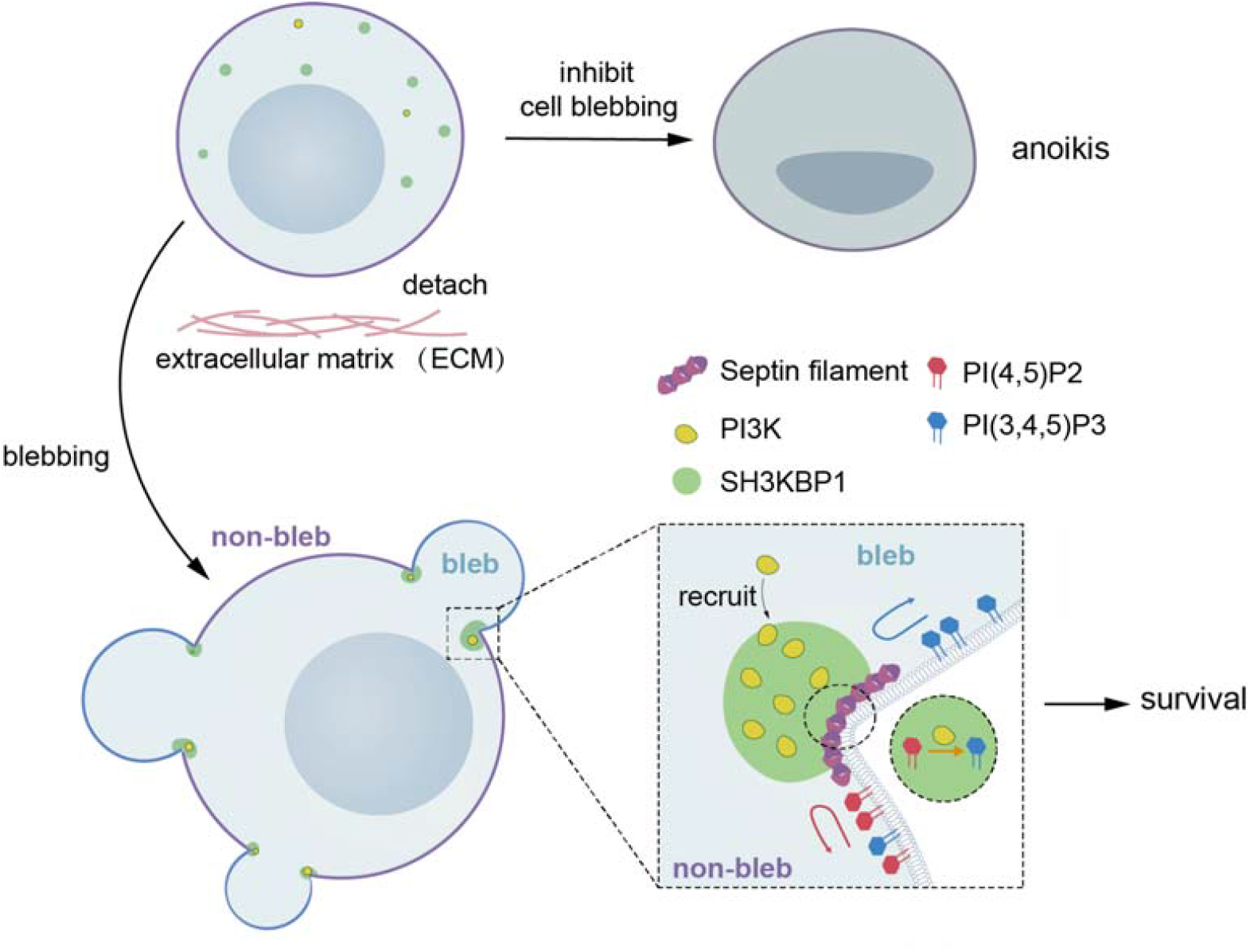

## Introduction

Adhesion-dependent survival is a fundamental property of cells in physiological processes. The sites where cells attach to the extracellular matrix or neighboring cells-such as focal adhesions and adherens junctions, can initiate signals that promote cell survival^1,2^. These structures act as central hubs for signaling molecules, protein scaffolds, and effector proteins in their vicinity. Without proper attachment, most cells undergo a programmed cell death process known as anoikis^3,4^. However, metastasizing tumor cells showed strong anoikis resistance when they break through the basement membrane and enter the vessels or abdominal cavity. Hence, acquiring molecular mechanisms that grant resistance to anoikis is a pivotal stage in the development of cancer^5,6^.

When detachment occurs, cells may become rounded and display dynamic surface blebs^7-10^. These blebs are hemispherical protrusions driven by internal hydrostatic pressure at locations where the actin cortex is disrupted^11^. While blebs are commonly observed during apoptosis, there exists a separate category termed as ‘dynamic blebs’ found in healthy cells with intact cortices, often linked to processes such as cellular detachment, mitosis, and amoeboid movement^12-14^.Non-malignant cells that become detached typically exhibit dynamic blebbing for a limited period of 1–2 hours before undergoing anoikis if they fail to re-establish attachment^4,9,15^. In contrast, cancer cells display high resistance to anoikis, enabling them to maintain rounded blebby shapes in environments with little or no attachment. These cells often adopt an amoeboid-like phenotype both in vivo and in vitro, notably at the invasive edge of tumors^16-20^. This enhanced blebbing has recently been linked to increased metastatic potential^16,21^, suggesting that the morphological characteristics may serve as an indicator of disease aggressiveness.

More recent work suggests that blebbiness may also convey a survival advantage ^20,22,23^. Formation of blebs results in the generation of micron-scale membrane curvature, including both negative intracellular curvature (in the bleb proper) and positive intracellular curvature (at the edge of blebs which were referred as bleb necks). Septins are the only known eukaryotic proteins that sense micron-scale positive curvature, and are recruited to bleb necks in detached cells, resulting in subsequent pro-survival signaling hub assembly^23-28^. Besides, septins can serve as a membrane diffusion barrier for subcellular compartmentalization^29-32^. Although it is reported that the protein composition of the bleb region is different from that of the surrounding, contiguous plasma membrane^33^, it has been less clear how the lipid composition of the blebs differs from that of other cellular membranes.

Phosphoinositides (PIs) are spatiotemporally enriched membrane lipids, and they are involved in regulating membrane dynamics and different types of cell death^34^. Specific kinases such as phosphoinositide 3-kinase (PI3K) mediate the conversion of phosphatidylinositol (4,5)-bisphosphate (PI(4,5)P2) to phosphatidylinositol (3,4,5) trisphosphate (PI(3,4,5)P3). In the process of blebbing, PI(3,4,5)P3 is shown to increase dynamic cell shape changes and protrusion formation, and conversely, PI(4,5)P2 content is associated with inhibited blebbing. However, there still lacks evidence of whether distinct phosphoinositides are present in the cell membrane and how they participate in the unique signaling functions of blebbing. In this study, we found the enrichment PI(3,4,5)P3 at bleb membrane as well as its restricted diffusion at bleb necks. We provide novel insights into the regulation of phosphoinositides dynamics within blebs and unveil the role of the Septin-SH3KBP1-PI3K complex in governing PIs distribution and diffusion at the bleb neck. Our findings shed light on the role of PIs signaling during cell blebbing in cancer cell behavior and metastasis and provide potential new therapy targets.

### Result1: PI(4,5)P2 and PI(3,4,5)P3 localize to distinct plasma membrane regions

Phospholipids intricately govern the biophysical properties of membranes. Aside from specific proteins involved in blebbing^23^, we asked whether there are specific features of phospholipids in cellular blebs. MDA-MB-231 cells were transfected with specific lipid biosensors and induced to form dynamic blebs using the actin polymerization inhibitor-Latrunculin B under cell detachment^11^. Intriguingly, unlike the uniformly distributed membrane marker-CAAX, specific PI(3,4,5)P3 sensor Akt-PH^35^ was enriched in blebs as quantified by a higher mean intensity on blebs than non-bleb plasma membrane (Fig. 1A). In contrast, the PI(4,5)P2 sensor PLCδ-PH^36^ showed robust residency on non-bleb plasma membrane (Fig. 1A). Quantification of the whole cell membrane fluorescence intensity reflect the distinct spatial distribution of PI(4,5)P2 and PI(3,4,5)P3 on bleb vs non-bleb regions. We then used a recently reported cell surface rendering method^50^ to process PI(4,5)P2 and PI(3,4,5)P3 biosensor signal and found the results consistent with 2D images, in which PI(4,5)P2 are primarily found in less blebby regions whereas PI(3,4,5)P3 are enriched in more blebby area (Fig. 1B). In contrast, the other membrane lipids like phosphatidylserine (PS) and cholesterol exhibited a homogeneous distribution similar to CAAX (Fig. 1B), indicating a specialized role of PIs in cell blebbing.

**Figure 1.**
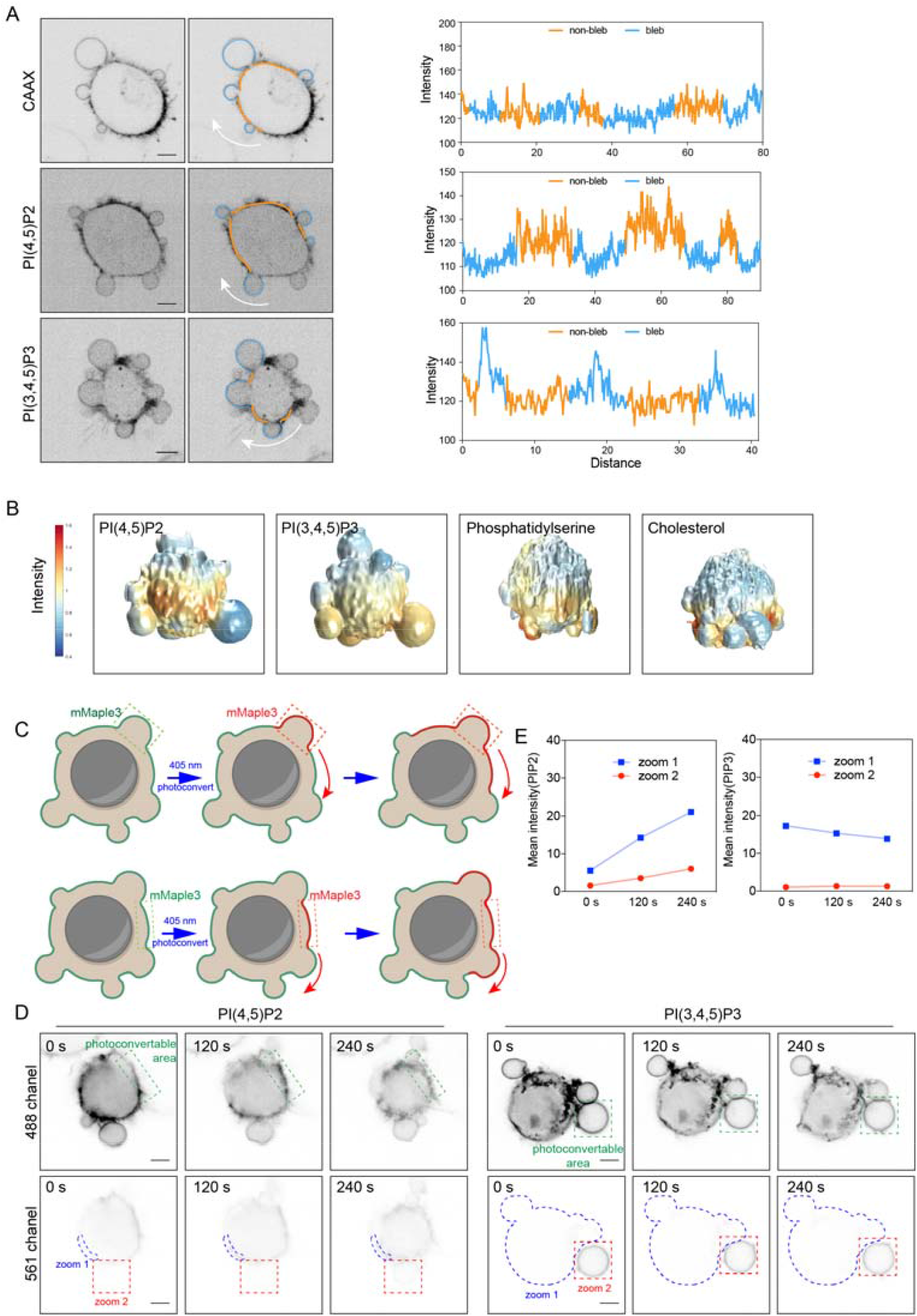
PI(4,5)P2 and PI(3,4,5)P3 exhibit distinct behaviors on different membrane compartments. (A)Left: representative images of mCherry-CAAX (upper), PI(4,5)P2 sensor PLCδ-PH-GFP (middle) and PI(3,4,5)P3 sensor Akt-PH-mCherry (bottom) in low concentration Latrunculin B (300 nM) treated MDA-MB-231 cell. Scale bar, 5 μm. Right: line scan plot showing the distribution of mCherry-CAAX (upper), PLCδ-PH-GFP (middle) and Akt-PH-mCherry (bottom) on cell membrane along the direction of the white arrowhead. (B)Surface renderings of intracellular PI(4,5)P2, PI(3,4,5)P3, phosphatidylserine and cholesterol mean intensity within 1Lµm of cell surface. (C)Schematic diagram of photoconvertible green fluorescent protein mMaple3 transition upon localized blue light activation and subsequent flow detection on cell membrane. (D)Time lapse images of MDA-MB-231 cells expressing PI(4,5)P2 sensor PLCδ-PH-mMaple3 (left) or PI(3,4,5)P3 sensor Akt-PH-mMaple3 (right) after transient photoconversion at specific membrane sites. Green dashed boxes are photoconverted sites. Blue and red dashed boxes marked sensor intensity are calculated in Fig. 1E. Scale bar, 5 μm. (E)Quantification of mean fluorescence intensity of dashed boxes in 1D. Left: PI(4,5)P2 mean intensity. Right: PI(3,4,5)P3 mean intensity. Zoom 1: non-bleb membrane. Zoom 2: bleb membrane.

To further dissect the mechanism underlying the unique distribution of PI(4,5)P2 and PI(3,4,5)P3, we built upon a photo-conversion system in which blue light-sensitive mMaple3 protein^37^ can switch from green signals to red as a mean to monitor the membrane lipid flow (Fig 1C). After converting a portion of PI(4,5)P2 sensor in non-bleb region using blue light, we found that the converted red signals could diffuse throughout the non-blebby membrane except for the blebby region. In contrast, the switched PI(3,4,5)P3 sensor signals were confined to the blebs, falied to diffuse into the surrounding non-blebby region. (Fig. 1D and E). The specific localization of PIs strongly suggests the involvement of phosphoinositide 3-kinase (PI3K) activity, which is responsible for converting PI(4,5)P2 to PI(3,4,5)P3. Furthermore, the inability of PI(3,4,5)P3 signals to flow out of the blebby regions indicates potential barriers to molecular diffusion or active mechanisms for retaining PI(3,4,5)P3 within blebs.

### Result2: PI3K colocalizes with Septin6 at bleb neck to mediate PIs distribution

PI3K is a key enzyme that converts PI(4,5)P2 to PI(3,4,5)P3, serving as a second messenger to activate downstream signaling pathways involved in cell survival, proliferation, and migration^38^.The distinct distribution of PI(4,5)P2 and PI(3,4,5)P3 at the cell bleb membrane raises an intriguing question about the potential involvement of PI3K in this cellular compartmentalization. By expressing PI3K regulatory subunit-PIK3R1, we found it colocalized with Septin6, which has been reported to be recruited to bleb neck (the border between the bleb and the rest of the membrane on the cell surface) ^23^ (Fig. 2A). This result indicates that PI3K and Septin6 may cooperate to modulate signaling events at this site. To determine if Septin localization at the bleb neck is necessary for PIK3R1 localization, we engineered a Septin2 mutant, Septin2 (33-306), which disrupts Septin polymerization as well as its localization at bleb necks^23^. First, Septin2 (33-306) abolished the specific localization of Septin at bleb neck as shown by the diffusive Septin6-GFP localization (Fig. 2B). Second, the colocalization of PIK3R1 with Septin6 at bleb necks was eliminated (Fig. 2B). Third, we observed that the enrichment of PI(3,4,5)P3 at the blebby region ceased, while PI(4,5)P2 demonstrated unhindered mobility through the blebs (Fig. 2C and D). In addition, PI3K inhibitor LY294002 treatment resulted in PI(3,4,5)P3 loss at bleb membrane and PI(4,5)P2 uniform distribution at whole membrane (Fig. 2E), indicating that both PI3K localization and activity at bleb necks are necessary for PI(3,4,5)P3 and PI(4,5)P2 distribution as well as their diffusion.

**Figure 2.**
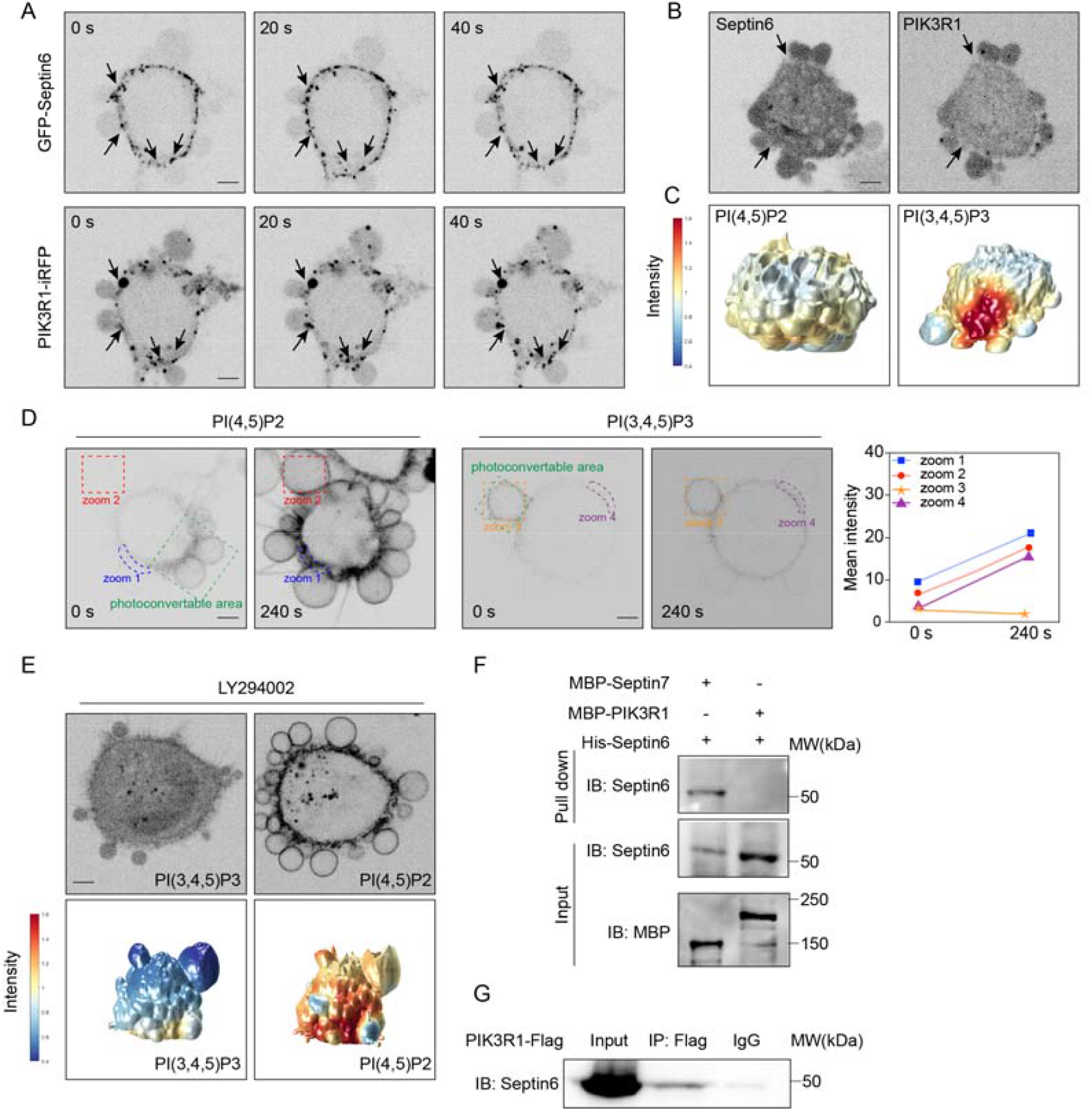
PIK3R1 resides at Septin6-enriched bleb neck and regulates PIs distribution. (A)Representative images of GFP-Septin6 (upper), PIK3R1-iRFP (bottom) in MDA-MB-231 cell during cell blebbing induced by low concentration Latrunculin B. Black arrows indicate Septin6 and PIK3R1 colocalization at bleb neck. Scale bar, 5 μm. (B)Representative images of GFP-Septin6 and PIK3R1-iRFP in a cell expressing SEPT2(33-306) polymerization mutant. Black arrows indicate bleb neck. (C)Surface renderings of intracellular PI(4,5)P2, PI(3,4,5)P3 mean intensity within 1Lµm of cell surface in cell expressing SEPT2(33-306) polymerization mutant. Scale bar, 5 μm. (D)Time lapse images of MDA-MB-231 cells expressing PI(4,5)P2 sensor PLCδ-PH-mMaple3 (left) or PI(3,4,5)P3 sensor Akt-PH-mMaple3 (middle) after transient photoconversion at specific membrane sites in cell expressing SEPT2(33-306) polymerization mutant. Green dashed boxes are photoconverted sites. Other colored dashed boxes marked sensor intensity are calculated in right panel. Scale bar, 5 μm. (E)Representative images and surface renderings of PI(4,5)P2 and PI(3,4,5)P3 intensity after treated with 10 µM LY294002 (PI3K inhibitor) 10 min. (F)MBP pulldown assay showing the interaction between Septin6 and Septin7 (on the left) or PIK3R1 (on the right). Samples were analyzed by SDS-PAGE and stained by Coomassie brilliant blue (N = 2 experiments, repeats are biological). (G)Co-immunoprecipitation assay showing interaction between PIK3R1-Flag and Septin6. (N = 3 experiments, repeats are biological).

However, GST-pull down assay indicated the absence of a direct interaction between PIK3R1 and Septin6 (Fig. 2F). In addition, co-immunoprecipitation validates the physical association between PIK3R1 and Septin6 (Fig. 2F). These results raise the possibility of scaffolding proteins serving as a physical link between PI3K and Septin6, facilitating functional cooperation despite the lack of direct binding at bleb neck.

### Result3: SH3KBP1 undergoes phase separation and bridges Septin6 and PI3K

We then sought to find out this link between Septin6 and PIK3R1. From the protein–protein interaction (PPI) networks of PIK3R1 and Septin6 which acquired through the STRING database, SH3KBP1 emerged as a potential mediator (Fig. 3A). We observed that SH3KBP1 exhibited strong co-localization with both Septin6 and PIK3R1 (Fig. 3B). Intriguingly, we observed that SH3KBP1 showed droplet-like shape both in the cytoplasm and at the bleb necks (Fig. 3B). Considering the morphological resemblance of SH3KBP1 with phase-separated non-membranous structures, we test whether it could undergo liquid-liquid phase separation (LLPS). Using an optogenetic approach based on the OptoDroplet system^39^, we found that CRY2-SH3KBP1 showed reversible formation of droplets both in the cytoplasm and at the bleb necks upon blue light exposure (Fig. 3C). In addition, SH3KBP1 droplets formation can recruit PIK3R1 into condensates (Fig. 3D).

**Figure 3.**
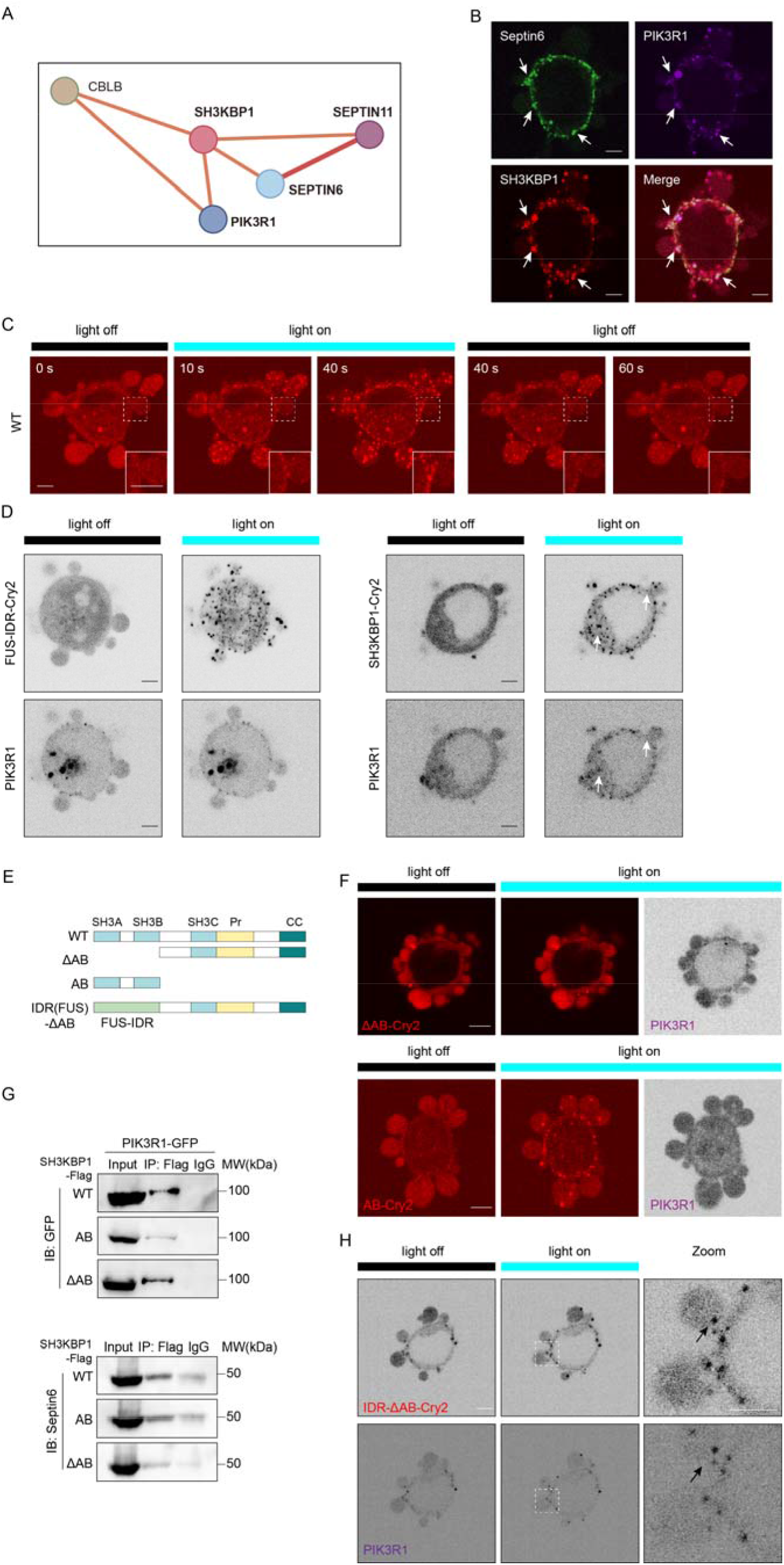
SH3KBP1 can phase separation and recruit PI3K regulatory subunit into condensates. (A)Protein–protein interaction (PPI) networks of PIK3R1 and Septin6 acquired through the STRING database(https://cn.string-db.org/). Thickness of the line indicate the degree of confidence prediction of the interaction.SH3KBP1 emerged as a potential linker between Septin6 and PIK3R1. (B)Representative images of GFP-Septin6 (upper left), PIK3R1-iRFP (upper right), SH3KBP1-mCherry-Cry2 (bottom left) and merged channel (bottom right) in MDA-MB-231 cell during cell blebbing induced by low concentration Latrunculin B. White arrows indicate Septin6-SH3KBP1-PIK3R1 colocalization at bleb neck. Scale bar, 5 μm. (C)Representative images of MDA-MB-231 cells expressing Opto-SH3KBP1 upon blue light exposure. Scale bar, 5 μm. Dashed boxes are zoomed in at the bottom right corner. Scale bar, 5 μm. (D)Left: representative images of MDA-MB-231 cells expressing Opto-FUS-IDR (upper) and PIK3R1-iRFP (bottom) upon blue light exposure. Right: representative images of MDA-MB-231 cells expressing Opto-SH3KBP1 (upper) and PIK3R1-iRFP (bottom) upon blue light exposure. White arrows indicate recruited PIK3R1 droplets by SH3KBP1. (E)Schematic representation of SH3KBP1 domain structure and ΔSH3-AB, SH3-AB, IDR(FUS)-ΔSH3-AB truncation design. (F)Representative images of Opto-ΔSH3-AB (upper) and Opto-SH3-AB (bottom) upon blue light exposure in PIK3R1-iRFP expressed MDA-MB-231 cells. Scale bar, 5 μm. (G)Left: co-immunoprecipitation assay showing interaction between PIK3R1-GFP and WT-SH3KBP1-Flag (upper) or SH3-AB-Flag (middle) or ΔSH3-AB-Flag (bottom) Right: co-immunoprecipitation assay showing interaction between Septin6 and WT-SH3KBP1-Flag (upper) or SH3-AB-Flag (middle) or ΔSH3-AB-Flag (bottom) (N = 3 experiments, repeats are biological). (H)Representative images of MDA-MB-231 cells expressing Opto-IDR(FUS)-ΔSH3-AB (upper) and PIK3R1-iRFP (bottom) upon blue light exposure. Scale bar, 5 μm. Dashed boxes are zoomed in (right). Scale bar, 5 μm.

SH3KBP1 contains three Src homology 3 (SH3) domains at the N-terminus, a proline-rich region in the center and a coiled-coil domain at the C-terminus^40^. Here we generated a set of SH3KBP1 truncations to figure out the function of different domains (Fig. 3E). We truncated SH3-A and SH3-B domains and found this truncation (ΔSH3-AB) lost the ability of phase separation (Fig.3F). Interestingly, ΔSH3-AB still maintained the interaction with Septin6 and PIK3R1 (Fig. 3G), but failed to recruit PIK3R1 into condensates (Fig. 3F),in contrast to full length SH3KBP1 (Fig. 3D). This result shows that the phase separation ability of SH3KBP1 is necessary for the enrichment of PIK3R1 and may further participate in mediating the specific distribution of phosphoinositides. Furthermore, we generated a mosaic protein (IDR (FUS)-ΔSH3-AB) by swapping the SH3-A and B domains of SH3KBP1 with intrinsically disordered region (IDR) in FUS protein^41,42^. IDR (FUS)-ΔSH3-AB robustly restored droplet formation property and recovered the PIK3R1 recruitment to bleb necks (Fig. 3H), while phase separated FUS-IDR droplets failed to do this (Fig. 3D), indicating PIK3R1 is specifically concentrated by SH3KBP1 droplets. In addition, we found SH3-AB truncation, which contains only SH3-A and SH3-B domains, could undergo phase separation. However, SH3-AB does not interact with PIK3R1 (Fig. 4B), indicating that PIK3R1 enrichment by SH3KBP1 not only depends on the phase separation but also the protein binding domains in SH3KBP1.

**Figure 4.**
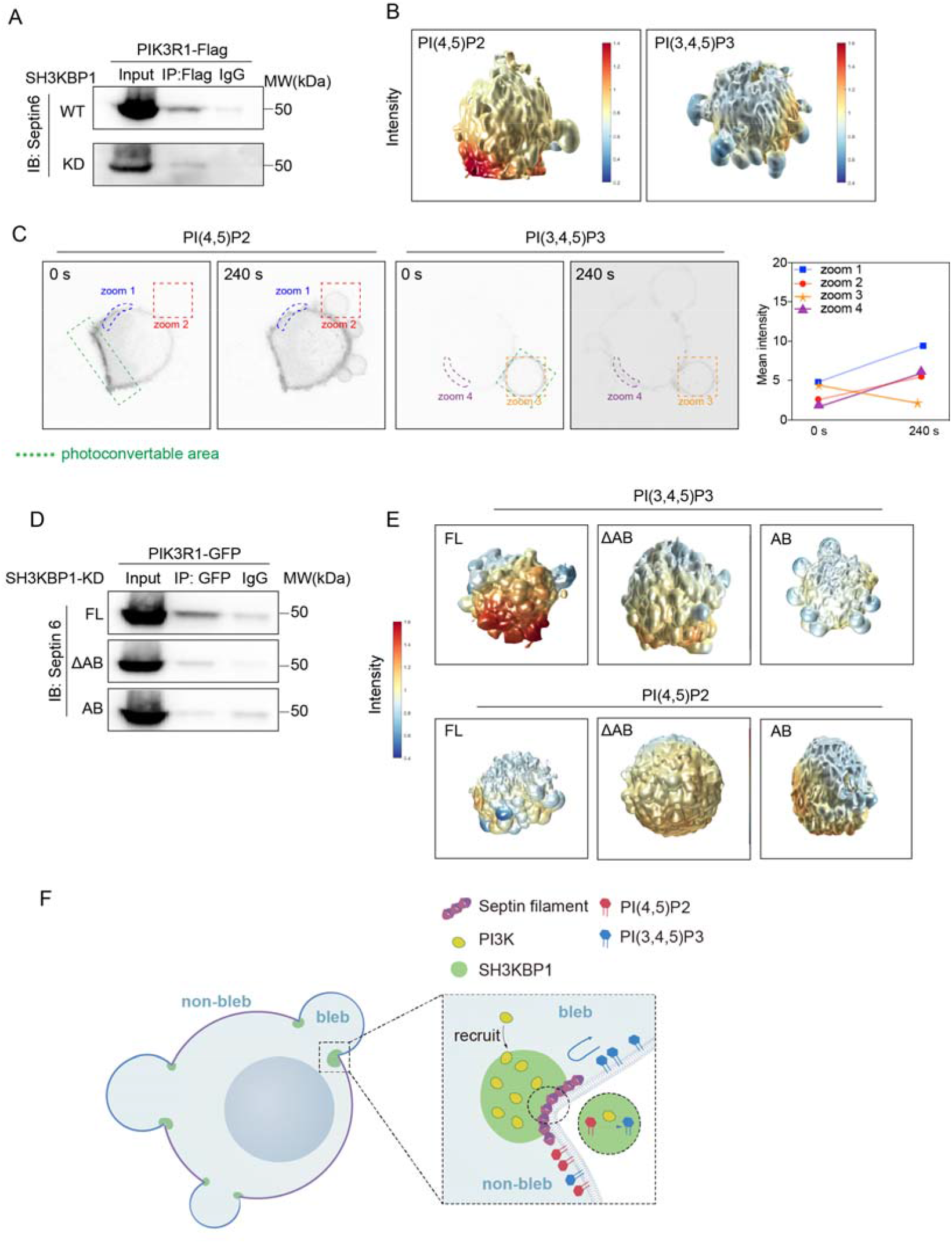
SH3KBP1 functions as the bridge for linking Septin and PIK3R1. (A)Co-immunoprecipitation assay showing interaction between PIK3R1-Flag and Septin6 in WT and SH3KBP1-KD MDA-MB-231 cells. (N = 3 experiments, repeats are biological). (B)Representative surface renderings of PI(4,5)P2 and PI(3,4,5)P3 intensity in SH3KBP1-KD cells. (C)Time lapse images of MDA-MB-231 cells expressing PI(4,5)P2 sensor PLCδ-PH-mMaple3 (left) or PI(3,4,5)P3 sensor Akt-PH-mMaple3 (middle) after transient photoconversion at specific membrane sites in SH3KBP1-KD cells. Green dashed boxes are photoconverted sites. Other colored dashed boxes marked sensor intensity are calculated in right panel. Scale bar, 5 μm. (D)Co-immunoprecipitation assay showing interaction between Septin6 and PIK3R1-GFP in SH3KBP1-KD cells after FL-SH3BKP1-Flag rescue (upper) or ΔSH3-AB-Flag (middle) or SH3-AB-Flag (bottom) (N = 3 experiments, repeats are biological). (E)Representative surface renderings of PI(4,5)P2 (bottom) and PI(3,4,5)P3 (upper) intensity in SH3KBP1-KD cells after FL-SH3BKP1-Flag rescue (left) or ΔSH3-AB-Flag (middle) or SH3-AB-Flag (right). (F)A proposed model for bleb regulated PIs behavior: Septin-SH3KBP1-PI3K condensates localize at bleb neck and regulate PI(4,5)P2 transition to PI(3,4,5)P3, resulting in distinct PIs distribution on plasma membrane.

Then, we sought to explore whether SH3KBP1 can bridge Septin6 and PIK3R1. SH3KBP1 knock-down (KD) disrupted the interaction between Septin6 and PIK3R1 (Fig. 4A), as well as the distinct distribution and residency of PI(3,4,5)P3 and PI(4,5)P2 (Fig. 4B and C). Upon reintroducing full length SH3KBP1 into KD cells, we rescued the interaction between Septin6 and PIK3R1 (Fig. 4D), along with a striking restoration of distinct PI(4,5)P2 and PI(3,4,5)P3 distribution on the blebbing cell membrane (Fig. 4E). However, the two truncations (ΔSH3-AB and SH3-AB) failed to rescue this phenotype (Fig.4D and E), indicating that both the phase-separation capacity of SH3KBP1 along with its interaction with PIK3R1 are crucial for the distribution of phosphoinositides (PIPs) in blebbing cells. Taken together, we raised a model wherein the Septin-SH3KBP1-PI3K axis localizes to bleb necks and regulates the distinct distribution of PI(3,4)P2 and PI(3,4,5)P3 on the bleb membrane (Fig. 4F).

### Result 4: Septin-SH3KBP1-PI3K complex collaboratively promotes cell survival under ECM detachment

During metastasis, malignant cells detach from the extracellular matrix and manifest dynamic surface blebs to resist anoikis^23^. Here, we investigated the role of blebbing in anoikis resistance by employing Latrunculin B (LatB) and wheat germ agglutinin (WGA) treatments to modulate bleb formation under detached conditions. Our findings revealed that the LatB treatment group exhibited a reduced level of cell death compared to the control group treated with DMSO, as evidenced by propidium iodide (PI) staining (Fig. 5A). Conversely, inhibition of bleb formation with WGA resulted in an elevated level of cell death (Fig. 5B), underscoring the significance of blebbing in conferring resistance against anoikis.

**Figure 5.**
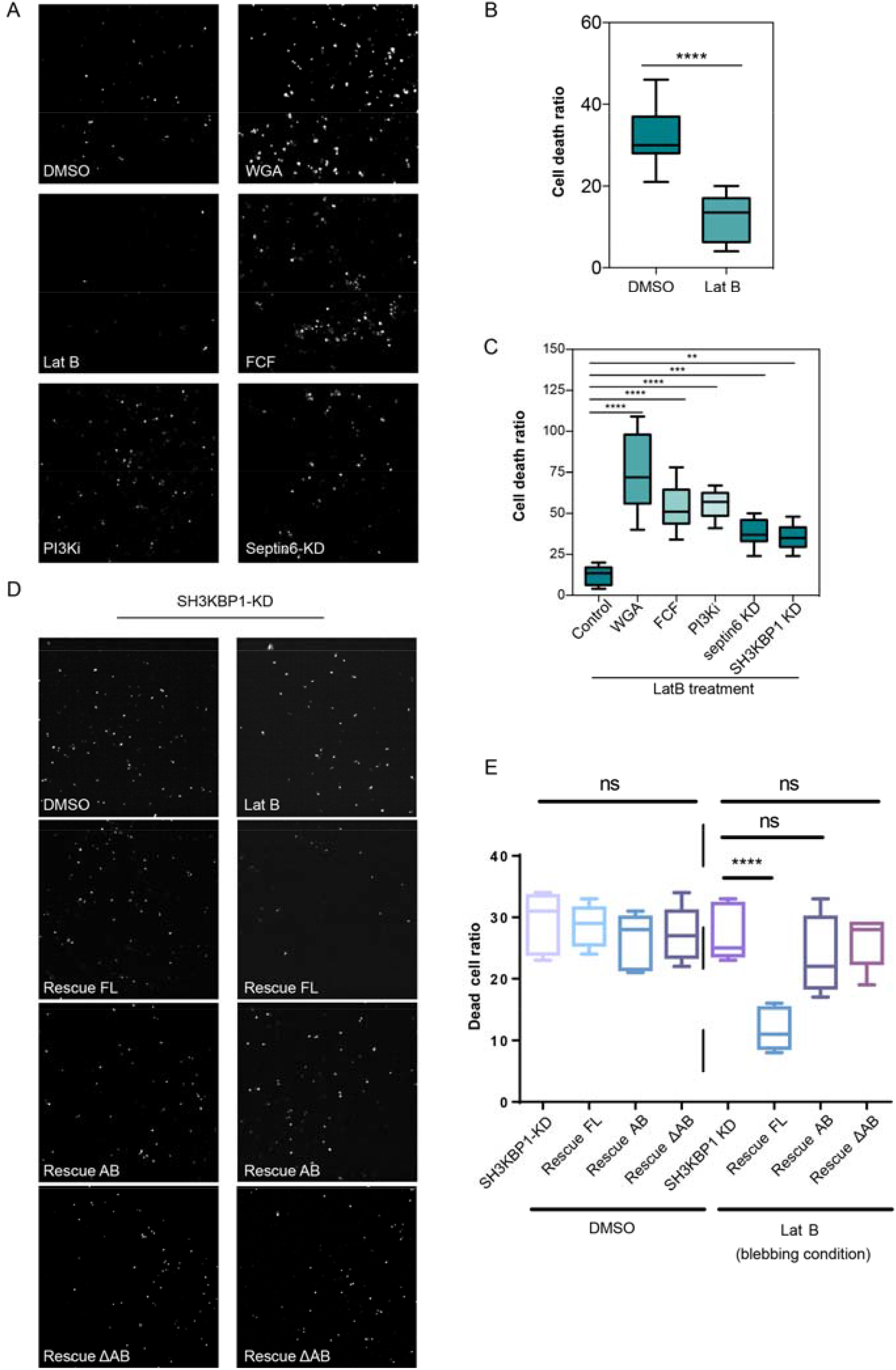
Septin-SH3KBP1-PI3K confer anoikis resistance in blebbing cells. (A)Representative PI staining images of cells under DMSO or LatB treatment cultured on substrates pre-treated with PLL-PEG after 24 h. Scale bar, 50 μm. (B)Bar chart shows the quantification of cell death ratio stained by PI in 5A (N_DMSO_ = 10, N_LatB_ = 10, error bar: mean with SEM, ****p < 0.0001 by unpaired Student’s t-test). (C)Representative PI staining images of cells under LatB treatment cultured on substrates pre-treated with PLL-PEG after treated with 10 μM WGA or 10 μM FCF or 10 μM LY294002 (PI3K inhibitor) 24 h. Scale bar, 50 μm. (D)Bar chart shows the quantification of cell death ratio stained by PI in Fig. 5C (N_Control_ = 10, N_WGA_ = 10, N_FCF_ = 10, N_PI3Ki_ = 10, N_Septin6 KD_ = 10, N_SH3KBP1 KD_ = 10, error bar: mean with SEM, ****p < 0.0001 by unpaired Student’s t-test, ***p < 0.01 by unpaired Student’s t-test, **p < 0.05 by unpaired Student’s t-test). (E)Representative PI staining images of SH3KBP1-KD cells rescued by FL, ΔSH3-AB or SH3-AB under DMSO (left) or LatB (right) treatment cultured on substrates pre-treated with PLL-PEG after 24 h. (F)Bar chart shows the quantification of cell death ratio stained by PI in 5E (N = 10, error bar: mean with SEM, ns_DMSO_ = 0.69333 by one-way ANOVA for multiple comparison, ****p< 0.0001, ns_SH3KBP1-KD versus Rescue AB_= 0.3099, ns_SH3KBP1-KD versus Rescue ΔAB_ = 0.7069 by unpaired Student’s t-test).

It has been reported that blebs recruit septins to cell surface and participate in anoikis resistance^23^, we then investigate whether the Septin-SH3KBP1-PI3K complex is involved in this process. To elucidate this, we evaluated the anoikis resistance of Septin6 knockdown cells or cells treated with the septin inhibitor forchlorfenuron (FCF). Consistent with our hypothesis, septin6 disruption markedly impaired anoikis resistance (Fig. 5A and C). Similarly, perturbation of PI3K activity or SH3KBP1 expression led to a significant decrease in cell survival (Fig. 5A and C), suggesting the involvement of these components in the anti-anoikis process.

Given SH3KBP1’s role as a signaling scaffold mediating interactions between Septin6 and PI3K, we further investigated its importance in resisting anoikis. By reintroducing full length SH3KBP1, as well as two SH3KBP1 truncations (ΔSH3-AB and SH3-AB), into SH3KBP1 knockdown cells, we observed that only full length SH3KBP1 could restore the rate of cell survival (Fig. 5D and E). These findings collectively underscore the crucial role of both the interaction between the Septin-SH3KBP1-PI3K complex and the phase separation ability of SH3KBP1 in facilitating anoikis resistance in blebbing cells.

## Discussion

Bleb formation has been widely acknowledged as a morphological program intricately linked with the progression of metastatic malignancies^12,16,17^. Recent study showed that dynamic blebbing prompts the construction of septin signaling hubs, which substitute for the loss of anchorage-dependent signaling activity caused by substrate detachment^23^. Aside from the protein hubs constructed during cell blebbing^23^, here we found PI(3,4,5)P3 and PI(4,5)P2 showed distinct distribution on bleb and non-bleb membrane and unidirectional flow during cell blebbing. We also identify that Septin-SH3KBP1-PI3K complex reside at bleb necks and regulating PIs distribution and diffusion in which SH3KBP1 bridge septin and PI3K through its phase separation and protein binding capacity.

PI(4,5)P2 and PI(3,4,5)P3 represent less than 1% of membrane phospholipids, however, they play an important role in numerous cellular processes^43,44^. PI(4,5)P2 is proposed to promote the interactions between plasma membrane and cytoskeleton, and bleb formation in cells can be a result of PI(4,5)P2 activity reduction at membrane^45^. PI(3,4,5)P3 is included in many plasma membrane-related processes^44^ and is important for bleb expansion as well as maintenance^16^. Here we speculate that the distinct distribution of PI(4,5)P2 and PI(3,4,5)P3 on bleb membrane could lead to persistent cell blebbing, thereby maintaining the signaling hubs at bleb necks for as long as possible to confer anoikis resistance.

Septins can polymerize into linear filaments, and then further assemble into higher-order structures, such as rings and gauzes^30^. Septins are also the only known eukaryotic proteins capable of detecting positive micrometer-scale membrane curvature^23^,^46^. Our results showed that septins polymerization disruption can result in PI(4,5)P2 and PI(3,4,5)P3 uniform distribution on non-bleb and bleb membrane. Although most research focus on the function of septin barrier for membrane proteins^30,47^, recent in vitro studies have shown that septins also reduce PI(4,5)P2 diffusion^48^. It raised the possibility that the physical barrier formed by septins influences the distribution of PI(4,5)P2 and PI(3,4,5)P3 as well as their dynamic flow between bleb and non-bleb regions.

Our experimental results reveal that the inhibition of SH3KBP1 leads to restored PI(3,4,5)P3 flow from bleb to non-bleb regions. In addition, we found SH3BKP1 can undergo phase separation and bridge septin and PI3K. It forms multimeric phase-separated droplets that recruit PI3K, facilitating their enrichment and anchoring at bleb neck sites, which raise the possibility that PI3K transverse PI(4,5)P2 to PI(3,4,5)P3 at bleb neck as a chemical barrier, creating PI(4,5)P2 and PI(3,4,5)P3 differential localization. Except for the SH3KBP1 droplets enrichment function, the phase separation induced droplets also have some physical properties. The wetting of the bleb neck membrane by SH3KBP1 condensates could potentially enhance lipid packing, promoting the formation of lipid microdomains at the neck^49^. This, in turn, may inhibit membrane fluidity in the neck region to regulate PIs flow.

Our model provides the possibility that the formation of Septin-SH3BKP1-PI3K complex at bleb necks contributes to the spatiotemporal regulation of cellular phosphoinositides distribution by creating a specialized membrane barrier with distinct physical and chemical properties. Understanding the complex interplay between bleb formation, phosphoinositide dynamics, and cytoskeletal organization holds immense therapeutic potential in the context of metastatic malignancies. Targeting key regulator identified in our study, SH3KBP1, may offer novel avenues for therapeutic intervention aimed at combating cancer invasion and metastasis.

### Limitations of the study

While this study provides significant insights into bleb promoted anti-anoikis, our study also has technical and conceptual limitations. First, the distinct distribution and diffusion patterns of PI(4,5)P2 and PI(3,4,5)P3 remain partially understood. The underlying mechanisms by which the Septin-SH3KBP1-PI3K complex regulates this unidirectional diffusion are still unclear. Second, it has been reported that septins can cause membrane compartmentalization by creating physical barriers or platforms that may affect the distribution and dynamic flow of PI(4,5)P2 and PI(3,4,5)P3 between bleb and non-bleb regions. However, the precise mechanism by which Septin-SH3KBP1-PI3K regulates PI(4,5)P2 and PI(3,4,5)P3 diffusion without influencing other lipids, such as phosphatidylserine (PS) and cholesterol, remains unknown. This is particularly noteworthy since cholesterol is a crucial lipid that regulates overall lipid diffusion. Third, our study did not employ single-molecule lipid tracking. Instead, we used a photo-conversion assay to detect the diffusion of PI(4,5)P2 and PI(3,4,5)P3. Although other studies have utilized single-molecule tracking to observe reduced PI(4,5)P2 diffusion in septin-enriched structures, this technique was not applied in our research, which may limit the precision of our findings regarding lipid diffusion. At last, although our data show that patients with SH3KBP1 overexpression have a reduced survival expectancy, it remains to be confirmed whether SH3KBP1 indeed undergoes phase separation under physiological or pathological conditions. Despite these limitations, our study advances the understanding of lipid diffusion and compartmentalization in blebbing membranes and provides a foundation for future research into the regulatory mechanisms of PI(4,5)P2 and PI(3,4,5)P3. Further studies addressing these limitations will be crucial for a more comprehensive understanding of lipid dynamics.

## Methods

### Cell Culture

MDA-MB-231 cells were generously provided by Yujie Sun laboratory (Peking University, China). Cells were cultured in Dulbecco’s modified Eagle medium (DMEM, Corning, 10-013-CRVC) supplemented with 10% fetal bovine serum (FBS, PAN-Biotech, P30-3302), 100 U mL-1 penicillin and 100 mg/mL streptomycin at 37 °C with 5% CO2. For cell passage, cells were washed with DPBS (Macgene, CC010) and digested with trypsin (Macgene, CC012). For detached cell culture, cells were grown to 70–80% confluency, trypsinized for 3 min, resuspended at approximately 100LcellsLml^-1^ (measured by eye using light microscopy). Drug treatments was added to suspensions if appropriate to the experiment, which were then immediately transferred to uncoated glass-bottom cell culture dishes. Detached cells were examined by light microscopy to confirm that cells did not aggregate-if aggregates were found, the experiment was discarded and repeated.

### Western Blot

For western blotting, cells were washed with DPBS once and lysed in an appropriate volume of RIPA buffer (50 mM Tris-HCl, pH 8.0, 150 mM NaCl, 1% Triton X-100, 0.5% Na-deoxycholate, 0.1% SDS, 1 mM EDTA and protease inhibitor cocktail) for 20 min on ice. Lysates were centrifuged at 13,572 g for 10 min and the supernatants were collected. Then, 5× SDS loading buffer was added to the supernatants and they were boiled for 10 min at 95°C. Protein samples were run on 10% SDS-PAGE acrylamide gels and transferred onto nitrocellulose (NC) membranes by wet electrophoretic transfer, followed by primary and second antibody incubation at 4°C overnight or room temperature for 2 h. Blots were visualized and recorded using SH-Compact 523 (SHST). The images were processed by ImageJ software.

### Live-cell imaging

For long-term live cell imaging, cells were plated on glass-bottom cell culture dishes and maintained in CO2-independent DMEM (Gibco, 18045-088) supplemented with 10% FBS, 100 U/mL penicillin and 100 μg/mL streptomycin at 37°C throughout the imaging process. For Opto-droplet phase separation imaging, dishes were covered with tin foil to block light and images were acquired at indicated intervals using a 63×1.4 NA objective lens on Andor Dragonfly confocal imaging system.

### Co-Immunoprecipitation

Cells were lysed with IP lysis buffer (25 mM Tris (pH 7.4), 1 mM EDTA, 150 mM NaCl, 5% glycerol, 1% NP□ 40, pH 7.4) for 15 min on ice. The cell lysis was centrifuged at 13572 g for 10 min at 4□ °C to remove the insoluble components. 80 µL of lysates were taken into a clean microfuge tube as “input” samples. The supernatants were added with antibodies (1 µg antibody per milligram protein) and rotated at 4□°C overnight. Then the protein A and G beads were washed with IP lysis buffer and added to the antibodiesLsupernatants mixtures followed by 3 h rotating at 4□°C. Finally, the beads were collected and washed three times with lysis buffer and boiled in 1x SDS sample buffer. 5x SDS loading buffer was used to boil the “input” samples. To detect the interaction proteins, the samples were separated by SDS-PAGE gels. To analyze protein interactome, the samples were separated by SDS-PAGE gels and then the target gels were collected for mass spectrum.

### Plasmid construction and DNA transfection

The plasmids construction used an ABclonal MultiF Seamless Assembly kit (ABclonal, RK21020). SH3KBP1-GFP, PLCδ-PH-GFP, GFP-Septin6, PIK3R1-GFP were subcloned into lentiviral vector (pLVX-AcGFP-N1) using EcoRI and BamHI enzymes from Transgene (JE201 □ 01, JB101 □ 01) for establishing AcGFP □ fused plasmids. And PIK3R1-iRFP were subcloned into pLVX-AcGFP-N1 with C-terminal tagged iRFP. For mCherry-Akt-PH, mCherry and EGFP fragments were amplified by regular PCR and then inserted into N-terminus of Akt-PH gene on pLVX-AcGFP-N1 vector using EcoRI and NotI enzymes (Transgene, JX101-01). Plasmid containing Cry2 was a gift from Chandra Tucker (Addgene plasmid # 60032, http://n2t.net/addgene:60032, RRID: Addgene_60032) to generate the Opto-droplet Optogenetic system. For PIK3R1-Flag, SH3KBP1-Flag, ΔSH3-AB-Flag, SH3-AB-Flag, Septin6-Flag, PIK3R1, SH3KBP1, ΔSH3-AB, SH3-AB or Septin6 was fused with a Flag tag contained in a short PCR primer into pLVX-AcGFP-N1 backbone.

For transient transfection, MDA-MB-231 cells seeded on 6-well plates were transfected with 2 μg midi-prep quality plasmid DNA in Opti-MEM (Invitrogen, 31985-070) containing 2 μL Neofect™ DNA transfection reagent (TF20121201) following the protocol, for 24-48 h. After that, cells were checked for fluorescent protein expression followed by other experiments. As for virus particle generation, HEK293T cells seeded on 10 cm culture dishes (Nest, 704201) were co-transfected with 5 μg of the expression plasmid, with 3.75 μg packaging plasmids psPAX2, and 1.25 μg pCMV-VSV-G, in Opti-MEM containing 10 μL Neofect™ DNA transfection reagent. To get a stable cell line, MDALMBL231 cells were infected 3 times with 1 mL viral supernatant, 1 mL DMEM supplemented with 2 µL polybrene and incubated at 37°C for 12–24 h.

### Reagents

The following reagents were used in this study: PI3K inhibitor LY294002 (Beyotime-S1737), actin polymerization inhibitor Latrunculin B (LatB, Sigma-Aldrich-L5288), Forchlorfenuron (FCF) from Sellck-S5306, Wheat Germ Agglutinin (WGA) from Bioss-D20067.

The following antibodies were used in this study: mouse anti-α-tubulin (T9026, 1:5000 for western blotting) from Sigma-Aldrich; rabbit anti-Septin6 (HPA005665, 1:4000 for western blotting) from Sigma-Aldrich, rabbit anti-SH3KBP1(Abclonal-A1952), mouse anti-GFP (M048-3, 1:4000 for western blotting and 1 uL antibody per milligram protein for Co-IP) from MBL International; anti-mouse (sc-516102, 1:4000) and anti-rabbit (sc-2004, 1:4000) horseradish peroxidase (HRP)-conjugated secondary antibodies from Santa Cruz Biotechnology.

### Anoikis Assay

Glass slides were coated with PLL-PEG (5 µg mL^-1^ in PBS) for 1 h at 37 °C to prevent cells from adhering to the base of culture dishes. This was performed to mimic the anchorageLindependent growth conditions of the cells.

### Microscopy imaging of cell death

Cell death was evaluated by propidium iodide (PI, Sigma-Aldrich, P4170-10MG) uptake of cells measured by fluorescence imaging. PI was added to the culture medium at a final concentration of 5 μg/mL before confinement. Static images of PI positive cells were captured using a Plan Apo 4x air objective lens on a Spinning Disk Confocal Microscope (Andor Dragonfly). Time-lapse imaging of cell death was imaged via epifluorescence microscopy (LIFE EVOS FL Auto fluorescent microscope, Life Technologies, USA). Raw full bit depth images were processed using ImageJ software (https://imagej.nih.gov/ij/). Image data shown were representative of at least three randomly selected fields.

### Confocal microscopy on cells expressing photoconvertible mMaple3

MDA-MB-231 cells were transfected with mMaple3-PLCδ-PH or Akt-PH construct 24 hours before and subjected to live cell imaging on Leica STELLARIS 8 microscope. mMaple3 can be switched from a green state to a red state upon activation with 406-nm light. Cells were imaged using 488 nm and 559 nm laser wavelengths to confirm emission of fluorescence in green channel but not red. Local complete photoconversion of green fluorescence to red was achieved by exposing cells to 406-nm light for 2Lseconds at 10% lamp power and confirmed by immediate cell imaging with 488 nm and 559 nm laser wavelengths. After photoconversion, cells were imaged in the red and green channel to visualize the diffusion. Image analysis was performed in ImageJ software.

### MBP Pull down

20 μg proteins with MBP tag (MBP-Septin7/MBP-PIK3R1) were first incubated with 20 μL dextrin beads slurry (Smart-Lifesciences, SA077010) in buffer containing 20 mM Tris (pH 8.0), 150 mM NaCl, 1% glycerol and 0.5 mM tris (2-carboxyethyl) phosphine hydrochloride (TCEP) at 4 °C for 3 h. Then dextrin beads were centrifuged, and the supernatants were discarded. After that, 20 μg proteins without MBP tag (His-Septin6) were added and incubated for another 30 min. After washing two times with buffer containing 20 mM Tris (pH 8.0), 500 mM NaCl, 1% glycerol and 0.5 mM TCEP, bound proteins were denatured with 2x SDS loading buffer, subjected to SDS–PAGE and stained by Coomassie brilliant blue /visualized on Tanon-5200 Chemiluminescent Imaging System (Tanon Science & Technology).

### Opto-droplet system

For Opto-droplet phase separation imaging, cells transfected with SH3KBP1-mch-Cry2 were covered with tin foil to block light and the droplets of SH3KBP1 can reversibly form upon blue light exposure. Images were acquired at indicated intervals using a 63×1.4 NA objective lens on Andor Dragonfly confocal imaging system.

### Fluorescence recovery after photobleaching (FRAP) assay

Cells were cultured in glass-bottom dishes at least 8 h before imaging. FRAP assay was performed on a ZEISS Laser Scanning Microscopes 880 at 37°C. GFP signals in regions of interest (ROI) were bleached using a 488-nm laser beam at full power. The fluorescence intensity between pre-bleaching and the time point right after bleaching was recorded by microscope. In order to extract quantitative information from the FRAP curves, logarithmic equation fitting, and linear equation fitting were used multiple times in Microsoft Excel.

### 3D cell image analysis

Cell morphology and PIs localization were analysed principally using u-shape3D method from Danuser article^23,53^. Most code can be found at https://github.com/DanuserLab.

### Statistics and Data Display

The number of biological and technical replicates and the number of samples are indicated in figure legends and the main text. Data are mean ± SEM, ± SD or 95% CI. as indicated in the figure legends and supplementary figure legends. Paired/unpaired Student’s t-test and one-way ANOVA were performed with GraphPad Prism 7.0 and Excel (Microsoft). Data from image analysis was graphed using Prism 7.0.

